# GDF15 Concentrations in Maternal Serum Associated with Vomiting in Pregnancy: the Cambridge Baby Growth Study

**DOI:** 10.1101/221267

**Authors:** Clive J. Petry, Ken K. Ong, Keith A. Burling, Peter Barker, John R.B. Perry, Carlo L. Acerini, Ieuan A. Hughes, David B. Dunger, Stephen O’Rahilly

## Abstract

Nausea and vomiting in pregnancy (NVP) affects 70-90% of all pregnant women but its pathogenesis is unknown. Growth and Differentiation Factor 15 (GDF15), secreted from the trophoblast and decidual stromal cells, is present at high levels in the blood of pregnant women. The receptor for GDF15 has recently been identified and is specifically expressed in the hindbrain where it transmits aversive signals including nausea and conditioned taste aversion. We explored the relationship between GDF15 concentrations in maternal serum during pregnancy and self-reported NVP. In a study of 791 women from the Cambridge Baby Growth Study maternal GDF15 concentrations were higher in women who reported vomiting in the 2^nd^ trimester (geometric mean: 11,670 pg/mL; 95% confidence interval: 11,056-12,318) and were even higher in the eleven women who reported taking anti-emetics during pregnancy (13,376 (10,821-16,535) compared to those who reported no nausea or vomiting during pregnancy (10,657 (10,121-11,222); P=0.02 and P=0.04, respectively, adjusted for gestational age at sampling and maternal BMI). In conclusion serum GDF15 concentrations early in the second trimester of pregnancy are significantly and positively associated with second trimester vomiting and with maternal anti-emetic use. In the context of the recently revealed biology of GDF15 this data suggests that antagonism of GDF15 may have some potential for therapeutic benefit in NVP.

## Background

Nausea and vomiting in pregnancy (NVP) affects 70-90% of all pregnant women. The most severe form of NVP, hyperemesis gravidarum (HG), leads to maternal dehydration and electrolyte imbalance and is the most common cause for hospital admission during early pregnancy [1]. Even though the majority of cases of NVP are mild or moderate, these conditions have substantial consequences for the mother’s quality of life [2], psychological morbidity [3], workplace productivity [4], and diet quality, characterised by high intakes of carbohydrates and added sugar [5]. Furthermore, NVP may have potential adverse effects on the developing fetus, as indicated by higher likelihood of low birth weight, preterm delivery, and small size at birth for gestational age in women with HG [6]. While effective pharmacological interventions are available, there are concerns regarding possible fetal teratogenicity of some agents [7].

Despite the high prevalence of NVP, its pathogenesis is poorly understood. Protective factors include: non-primiparous pregnancy, older maternal age and smoking [1]. Reproductive hormones, such as human chorionic gonadotropin, progesterone and oestrogen, have been implicated due to their rise in concentrations in the mother’s circulation contemporaneous with the manifestation of NVP [8]. However nausea and vomiting are not common side-effects of such agents when administered in other settings. Recent genetic studies of HG have implicated rare variants in *TSHR*, which encodes the thyrotropin receptor [9], and *RYR2*, which encodes a stress-induced intracellular calcium release channel involved in cyclic-vomiting syndrome [10] in some familial cases. In contrast to rare and deleterious mutations as causes of HG, evolutionary theories have been proposed for NVP as a beneficial strategy to protect the fetus from maternal ingestion of noxious substances, particularly during the early stages of pregnancy coinciding with organogenesis when the fetus is most vulnerable [8].

Growth and Differentiation Factor 15 (GDF15) signalling through its receptor (a heterodimer of GFRAL and RET) has recently been identified to activate the mammalian chemoreceptor trigger zone of the medulla to suppress food intake in mice [11-14], and therefore represents a potential mechanism for the aversion to foods and eating behaviours during periods of stress, sickness or high vulnerability to external toxins [8]. GDF15 is expressed predominantly in liver, but also by other tissues including placental trophoblasts and decidua and circulating levels rise rapidly in maternal blood during the first trimester of pregnancy and remain elevated until delivery [15].

To explore the hypothesis that GDF15-GFRAL signalling may represent a fetal-induced system to promote protective aversive eating behaviours by the mother during early pregnancy [16], we measured circulating GDF15 concentrations in maternal serum during the 1^st^ half of pregnancy and related these levels to self-reported NVP in a large prospective pregnancy cohort study.

## Materials & Methods

### Cohort

The prospective Cambridge Baby Growth Study recruited 2,229 mothers (and their partners and offspring) attending antenatal ultrasound clinics during early pregnancy at the Rosie Maternity Hospital, Cambridge, United Kingdom, between 2001 and 2009 [17]. All mothers were over 16 years of age. In this cohort, 96.9 % of the offspring were of white ethnicity, 0.8 % were of mixed race, 0.6 % were black (African or Caribbean), 0.8 % were East-Asian, and 0.9 % were Indo-Asian. Blood samples, from which serum was separated and aliquoted, were collected from 1,177 (52.8%) mothers at recruitment.

Each mother was given a printed questionnaire at recruitment to fill in and return after the birth of their child [18]. The questionnaire included boxes to tick if the participants had experienced NVP during pregnancy. If either the nausea or vomiting boxes was ticked there were further boxes to complete concerning the timing (i.e. week(s) of pregnancy) when the nausea or vomiting was experienced. An additional question asked “Have you taken any medicine during this pregnancy?” and a table was provided for positive responses with the following headings: “Name”, “Disease”, “Daily Dose”, “No. of Days” and “Gestational Week(s)”. A total of 1,238 women (54.6 %) returned a questionnaire. Of these, only 3 self-reported that they had HG and a further 17 reported treatment with an anti-emetic agent: cyclizine (n=7), promethazine (n=5), prochlorperazine (n=4), metoclopramide (n=2), domperidone (n=2), prednisolone (n=2), chlorphenamine (n=1), ondansetron (n=1), chlorpromazine (n=1) and unknown (n=1). The timing of NVP was categorised into trimesters (1^st^ : up to gestational week 12; 2^nd^ 13 to 27 weeks; 3^rd^ 28+ weeks).

The current analysis was based on 791 women in CBGS who had an available serum sample collected between gestational age 12 and 18 weeks and returned a completed questionnaire [18].

### GDF15 Concentration Measurements

Serum GDF15 concentrations were measured using an in-house Meso Scale Discovery electrochemiluminescence immunoassay (Meso Scale Diagnostics, Rockville, Maryland, U.S.A.) developed using antibodies from R & D Systems Ltd. Quantikine reagents (BioTechne Ltd., Abingdon, U.K.). The sensitivity of this assay was 3 pg/mL and the working range went up to 32,000 pg/mL. Batch-to-batch variability was 9.8% at 352 pg/mL, 8.1% at 1490 pg/mL and 7.8% at 6667 pg/mL.

### Statistical Analysis

Women were categorised into one of three groups: vomiting; nausea but no vomiting; and no nausea or vomiting. The primary outcome was vomiting during the 2^nd^ trimester, as this coincided with the timing of maternal serum sampling. These were compared to women who reported no nausea or vomiting. Maternal body mass index (BMI) was calculated dividing self-reported body weight prior to pregnancy by self-reported height-squared.

Serum GDF15 concentrations were natural logarithm-transformed to achieve a normal distribution and were considered as the dependent variable in linear regression models with adjustment for gestational age at serum sample collection. Where the relationship with adjusted log-transformed GDF15 concentrations did not appear to be linear, data were transformed to approximate linearity prior to analysis. Statistical analyses were performed using Stata 13.1 (StataCorp LP, College Station, Texas, U.S.A.). P<0.05 was considered to indicate statistically significance.

## Results

37.7% (n=298) of women reported vomiting during any trimester of pregnancy. A further 37.9% (n=300) reported nausea but no vomiting, and only 24.4% (n=193) reported no nausea or vomiting. More women (32.0%, n=253) reported vomiting during the 1^st^ trimester compared to 22.1% (n=175) in the 2^nd^ trimester, and only 3.8% (n=30) in the 3^rd^ trimester. 86.9% and 56.7% of those reporting vomiting in the 2^nd^ and 3^rd^ trimesters also reported vomiting during the 1^st^ trimester, respectively. Women who reported vomiting during the 2^nd^ trimester had higher parity, were younger and were carrying relatively more female babies than women who reported no nausea or vomiting during pregnancy (Table 1), but there were no differences in BMI.

**Table 1.**
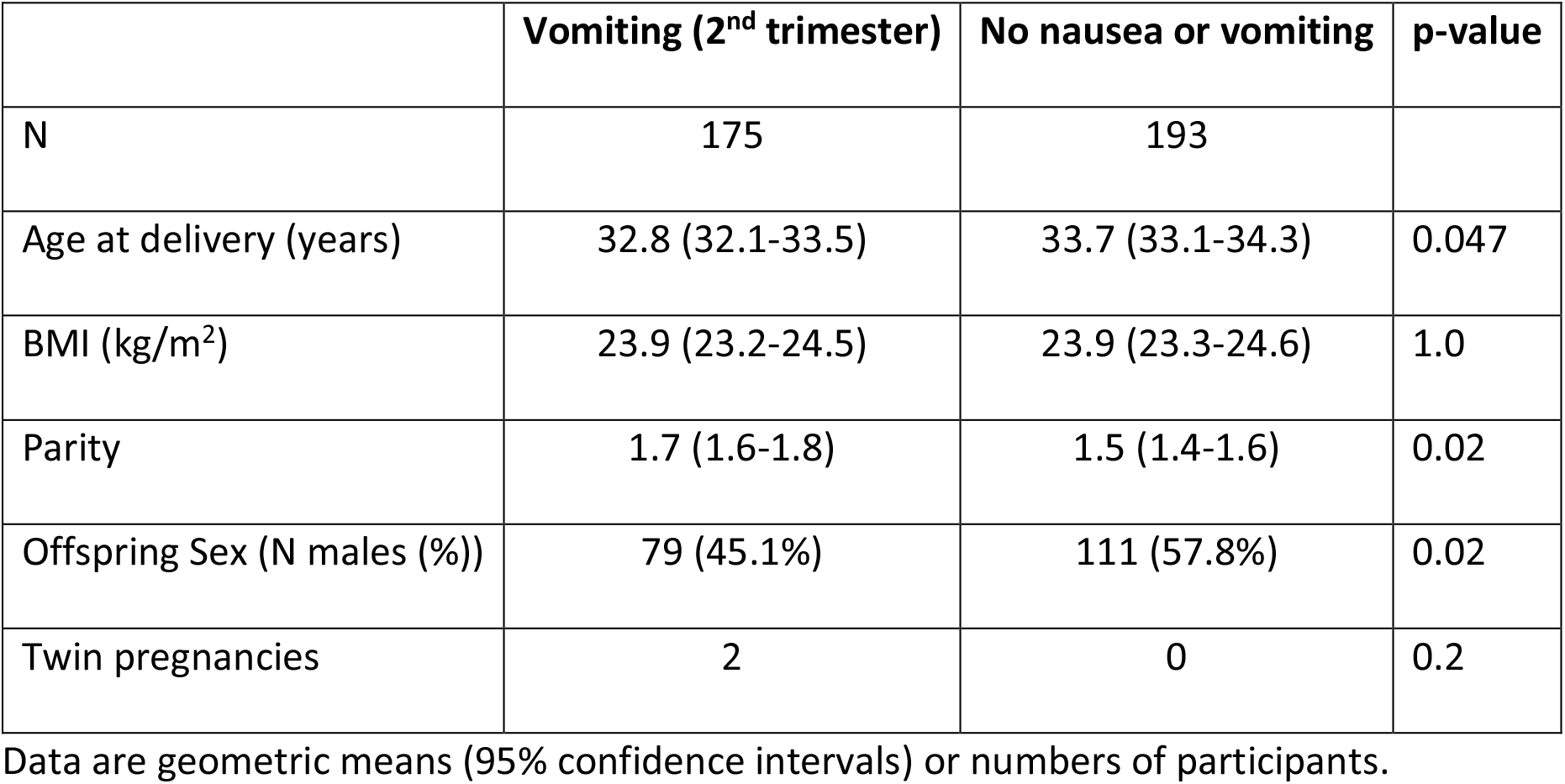
Clinical characteristics of the women who reported vomiting during the 2^nd^ trimester of pregnancy compared to women who reported no nausea or vomiting during pregnancy.

### Maternal GDF15 Concentrations

The median GDF15 concentration was 11,004 pg/mL (range 2,378-34,621) in serum samples collected at mean gestational age 15.1 weeks (range 12.0-18.0). Maternal GDF15 concentrations were not associated with gestational age at sampling (linear model with log-GDF15: P=0.4, standardised β=-0.03).

### Maternal GDF15 Concentrations Associated with Vomiting in Pregnancy

Maternal GDF15 concentrations were higher in women who reported vomiting in the 2nd trimester of pregnancy compared to those who reported no nausea or vomiting during pregnancy (P=0.02; Table 2). This association was unaltered by adjustment for gestation at serum sampling or by maternal BMI. The maternal GDF15 concentrations were unchanged in women who reported nausea alone in the 2nd trimester of pregnancy.

**Table 2.** Maternal GDF15 concentrations by self-reported vomiting in the 2^nd^ trimester or anti-emetic use during pregnancy.

| Group | n | Serum GDF-15 pg/mL | Unadjusted | Adjusted for gestational age | Additionally adjusted for maternal BMI |
| --- | --- | --- | --- | --- | --- |
| No nausea or vomiting | 193 | 10,593 (10,066-11,147) | Ref | Ref | Ref |
| Vomiting (2<sup>nd</sup> trimester) | 175 | 11,581 (10,977-12,219) | P=0.02 | P=0.02 | P=0.02 |
| Nausea without vomiting (2<sup>nd</sup> trimester) | 325 | 10,772 (10,328-11,235) | P=0.6 | P=0.6 | P=0.5 |
| Anti-emetic use (any trimester) | 11 | 13,157 (10,558-16,394) | P=0.06 | P=0.04 | P=0.04 |

Data are geometric means (95% confidence intervals).

Eleven women (1.4%) reported taking an anti-emetic medication during pregnancy (10 (90.9%) of whom reported vomiting, 1 (9.1%) of reported nausea without vomiting). Their serum GDF15 concentrations were also raised compared to women who reported no nausea or vomiting during pregnancy (P=0.04, adjusted for gestation; Table 2).

## Discussion

In this large prospective pregnancy cohort study, we show that maternal circulating GDF15 concentrations were higher in women who reported vomiting in the 2^nd^ trimester and were even higher in women who reported taking anti-emetics during pregnancy, compared to those who reported no nausea or vomiting during pregnancy.

To our knowledge, this is the first report relating GDF15 concentrations to vomiting during pregnancy. Circulating GDF15 concentrations rise rapidly in maternal blood during early pregnancy and several studies have reportedly substantially lower concentrations at around 6-13 weeks gestation in those pregnancies that subsequently miscarried [19, 20]. Possible explanations for this highly reproducible phenomenon have included the suggestion that maternal circulating GDF15 is a biomarker of successful placentation. Alternatively it has been suggested that GDF15 (also known as Macrophage inhibitory cytokine 1) may promote fetal viability through an immunomodulatory action. However the recent discovery of the highly specific expression of the receptor for GDF15 in the hindbrain makes this less likely [21]. Despite that uncertainty, our findings provide a possible mechanistic explanation for the widely observed associations between NVP and lower rates of miscarriage, malformations and prematurity [22].

Ideally, we would have measured maternal GDF15 concentrations in samples collected at gestational age 9 weeks, which is the peak for NVP symptoms. However, many women have not yet presented to maternal health services at that stage, and indeed for many women NVP represents the first indication of pregnancy. Maternal BMI was only available pre-pregnancy, but weight is unlikely to have changed much during the initial 15 weeks of pregnancy. Unsurprisingly, reflecting recruitment of the cohort at routine antenatal clinics, cases of HG were under-represented. Future case-control study designs are needed to test whether our findings can be extrapolated to HG.

There is a pressing need for improved therapy for NVP and HG. Manipulation of GDF15 and/or its signalling pathways may provide a novel avenue in this regard. Future studies are also needed to understand the role of fetal genetic variation and other factors that regulate maternal GDF15 concentrations during pregnancy.

It has been proposed that the role of GDF15 in the adult organism is to provide a signal to the brain that the organism is engaging in a behaviour that is damaging. Its hindbrain-localised receptor activates a highly aversive signal which promotes the future avoidance of this particular behaviour. We propose that the placenta has evolved to use the GDF15 system to promote food-aversive behaviours in the mother to protect the fetus from exposure to maternal ingestion of potential teratogens during the vulnerable stages of organ development.

## Declarations

### Ethics approval and consent to participate

The Cambridge Baby Growth Study was approved by the Cambridge Research Ethics Committee, Cambridge, United Kingdom. All procedures followed were in accordance with Good Clinical Practice guidelines. Written informed consent was obtained from all the study participants.

### Authors’ contributions

SOR hypothesised the involvement of GDF15 in NVP and initiated the project, which was designed by SO, DD, JP and KO. KAB and PB performed the serum measurements. CP collated Cambridge Baby Growth Study questionnaire data, prepared the serum samples and performed the statistical analyses. IH, DD, CA and KO designed and oversee the Cambridge Baby Growth Study. CP and KO drafted the manuscript, and all authors commented on subsequent drafts and approved the final manuscript.

## Acknowledgements

We thank all the families who took part in the Cambridge Baby Growth Study, and we acknowledge the crucial role played by the research nurses especially Suzanne Smith, Ann-Marie Wardell and Karen Forbes, staff at the Addenbrooke’s Wellcome Trust Clinical Research Facility, and midwives at the Rosie Maternity Hospital. We also thank Diane Wingate for expert assistance in sample preparation and Ashley Clarke for developing and validating the GDF15 immunoassay.

